# Monoclonal IgM antibodies raised against *Candida albicans* Hyr1 provide cross-kingdom protection against Gram negative bacteria

**DOI:** 10.1101/687442

**Authors:** Eman G. Youssef, Sondus Alkhazraji, Teclegiorgis Ghebremariam, Lina Zhang, Shakti Singh, Nannette Y. Yount, Michael R. Yeaman, Priya Uppuluri, Ashraf S. Ibrahim

## Abstract

Recent years have seen an unprecedented rise in the incidence of multidrug resistant (MDR) Gram negative bacteria (GNB) such as *Acinetobacter* and *Klebsiella* species. In view of the shortage of novel drugs in the pipeline, alternative strategies to prevent and treat infections by GNB are urgently needed. Previously, we have reported that the *C. albicans* hyphal-regulated protein Hyr1 shares striking 3D structural homology with cell surface proteins of *A. baumannii*; and active or passive vaccination with rHyr1p-N or anti-Hyr1p polyclonal antibody, respectively; protect mice from *Acinetobacter* infections. Here, we show that monoclonal antibodies (mAb) generated against Hyr1p, bind to the surface of *Acinetobacter* as well as *K. pneumoniae*. The anti-Hyr1 mAb also block damage to primary endothelial cells by the bacteria, and protect mice from lethal pulmonary infections mediated by *A. baumannii* and *K. pneumoniae*. Our current studies emphasize the potential of harnessing Hyr1p mAb as a cross-kingdom immunotherapeutic strategy against MDR GNB.

## Introduction

Infections caused by multidrug resistant organisms (MDRO) continue to pose a therapeutic challenge. In the past decade, *A. baumannii* has emerged as one of the most common MDRO of hospital acquired infections, causing a range of diseases from pneumonia to severe blood or wound infections ^1-6^. Of concern is that 40-70% of *A. baumannii* isolates are now extensively drug resistant (XDR; i.e. resistant to all antibiotics except colistin and tigecycline), reflecting a >15-fold increase since 2000 ^1,6-8^. Likewise, the *Enterobaceriaceae* organism *K. pneumoniae* causes high rates of morbidity and mortality in critically-ill, hospitalized patients, and in recent years have developed antimicrobial resistance to almost all classes of antibacterial drugs including carbapenamases ^9-11^. Together, *Acinetobacter* and carbapenam-resistant *K. pneumoniae* (KPC) have been flagged by the CDC as two of the top “Serious Threat Level Pathogens” due to resistance, failure of current standard of treatment, and high mortality rates. To make matters worse, the existing drug development pipeline against these bugs is unacceptably lean and it is almost certain that the organisms will develop resistance to any new approved antibiotics. Hence, novel strategies to prevent and treat life-threatening infections by the two species are urgently needed.

We have previously exploited innovative computational molecular modeling and bioinformatic strategies to discover novel vaccine and immunotherapy candidates that target more than one high priority pathogen. Our immunotherapeutics-discovery campaign culminated in the identification of *Candida albicans* Hyr1p, a <u>hy</u>phae <u>r</u>egulated cell surface protein that helps the fungus to resist phagocyte killing. Mice vaccinated with Hyr1p are protected from *C. albicans* infections ^12,13^. Recently, we found that the Hyr1p protein shares striking 3D structural, and immunological homologies to antigens present on the Gram negative bacteria (GNB) *A. baumannii*, including with the putative hemagglutinin/hemolysin protein FhaB, and a number of siderophore-binding proteins^14^. Polyclonal antibodies (pAbs) against peptides derived from the Hyr1p N-terminus blocked *A. baumannii-*mediated lung epithelial cell invasion, and killed the bacterium *in vitro* ^14^. Importantly, anti-Hyr1p pAb completely protected mice from *A. baumannii* infections. These results provided proof of concept for targeting Hyr1p for developing immunotherapies against GNB, and laid a groundwork for generation and evaluation of the efficacy of anti-Hyr1p monoclonal antibodies against MDR GNB.

In the current study, we generated monoclonal antibodies (mAb) against Hyr1p and show that these mAb not only recognize different clinical isolates of *A. baumannii*, but also bind to drug resistant *K. pneumoniae*. We further demonstrate the efficacy of these targeted mAb to block bacterial invasion to host cells, and protect mice against lethal pulmonary infection by both MDR bacteria. Given the alarming rate at which MDRO’s are growing as a global threat, a passive vaccination strategy using therapeutic mAbs serves as a highly desirable strategy to combat these difficult to treat infections either as a standalone or adjunctive therapies to antibiotics.

## Results

### Anti-HYR1 mAbs bind to Gram negative bacteria

We have previously reported that pAb raised against a 14 amino acid peptide of Hyr1p (LKNAVTYDGPVPNN; also called peptide#5) blocked virulence traits of *A. baumannii in vitro*, and completely protected diabetic and neutropenic mice from *Acinetobacter* bacteremia and pulmonary infection ^*14*^. Encouraged by these results, and to enhance the therapeutic potential of the antibodies, we developed monoclonal antibodies (mAb) against the same surface-exposed and immunodominant peptide. The mAbs (all produced antibodies belonged to the IgM isotype) were tested for their binding abilities to *C. albicans* and also the GNB *A. baumannii*, and *K. pneumoniae.* Four individual FITC labeled mAb clones (H1, H2, H3 and H4) at 100 *µ*g/ml were tested against three MDR GNB: *K. pneumoniae*-RM (KPC-RM, carbapenam-resistant isolate), *K. pneumoniae*-QR (KP-QR, MDR strain sensitive to carbapenam), and MDR *A. baumannii* (HUMC-1), and the extent of mAb binding to the bacterial surface was quantified by flow cytometry. The results were compared to isotype matching nonspecific control antibodies. Out of the four clones of mAb tested, H3 and H4 displayed the highest levels of binding to all tested GNB with at least 30 to 300-fold increase compared to the isotype matching control IgM (**Fig 1A**). Binding potential was also visualized by a shift in the peaks of the anti-Hyr1 IgM binding versus the isotype matching control antibodies (**Fig 1B**). The right shift in the peaks of individual mAb also correlated with their respective increase in mean fluorescence of the cells.

**Figure 1:**
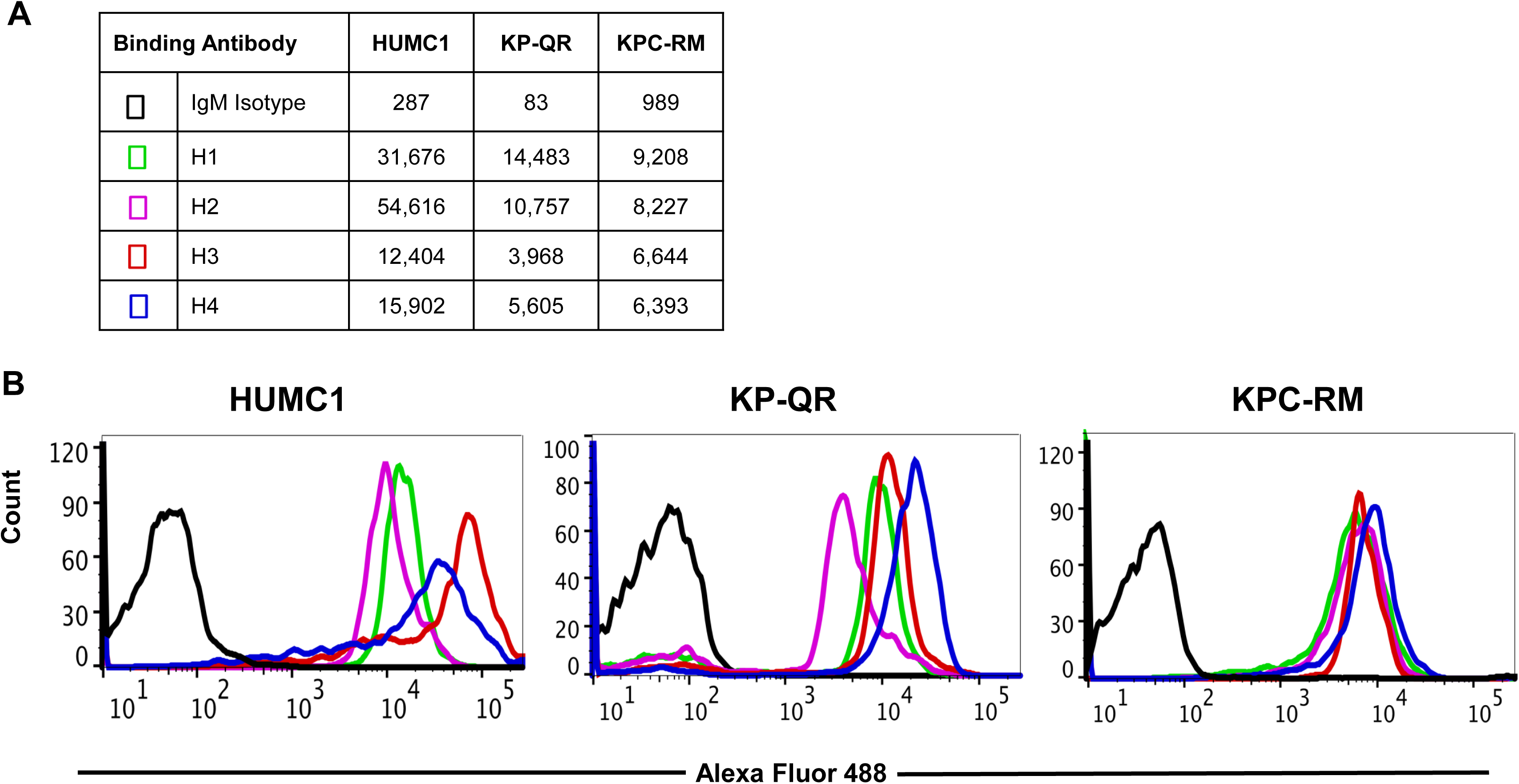
Binding of mAb targeting Hyr1p peptide 5 to Gram negative bacteria (GNB): mAb clones (and isotype-matched control IgM) were evaluated for binding to *A. baumannii* (HUMC-1), *K. pneumoniae*-QR (KP-QR) and *K. pneumoniae*-RM (KP-RM) at a concentration of at 100 *µ*g/ml. The extent of binding was quantified by flow cytometry after staining the bound antibodies with alexa 488-conjugated secondary antibody. Data was represented as mean fluorescence intensity of the Ab-bound bacteria. (**A**). The degree of binding was also visualized by a shift in the peaks in the anti-Hyr1 IgM binding conditions versus the control antibodies (**B**).

Next, we tested the sensitivity of the clones to recognize the GNB. Clone H3 bound to the three organisms even at low mAbs concentrations. Specifically, 30 *µ*g/ml of H3 displayed 50% of binding to both the KP-QR and *A. baumannii* MDR strain HUMC1, while 3 *µ*g/ml of the mAb bound to at least 10% of the total cells, which was significantly higher than the isotype-matching control IgM (**Fig 2**). We further evaluated the binding affinity of the two clones H3 and H4 against other drug resistant clinical isolates of *Acinetobacter* and *K. pneumoniae* (KPC). The mAbs bound KPC-6, KPC-8, HUMC-6 and HUMC-12 at significantly high affinity compared to the control IgM (**S1**). These results indicate that binding of mAbs to surface of the tested GNB is not isolate specific.

**Figure 2:**
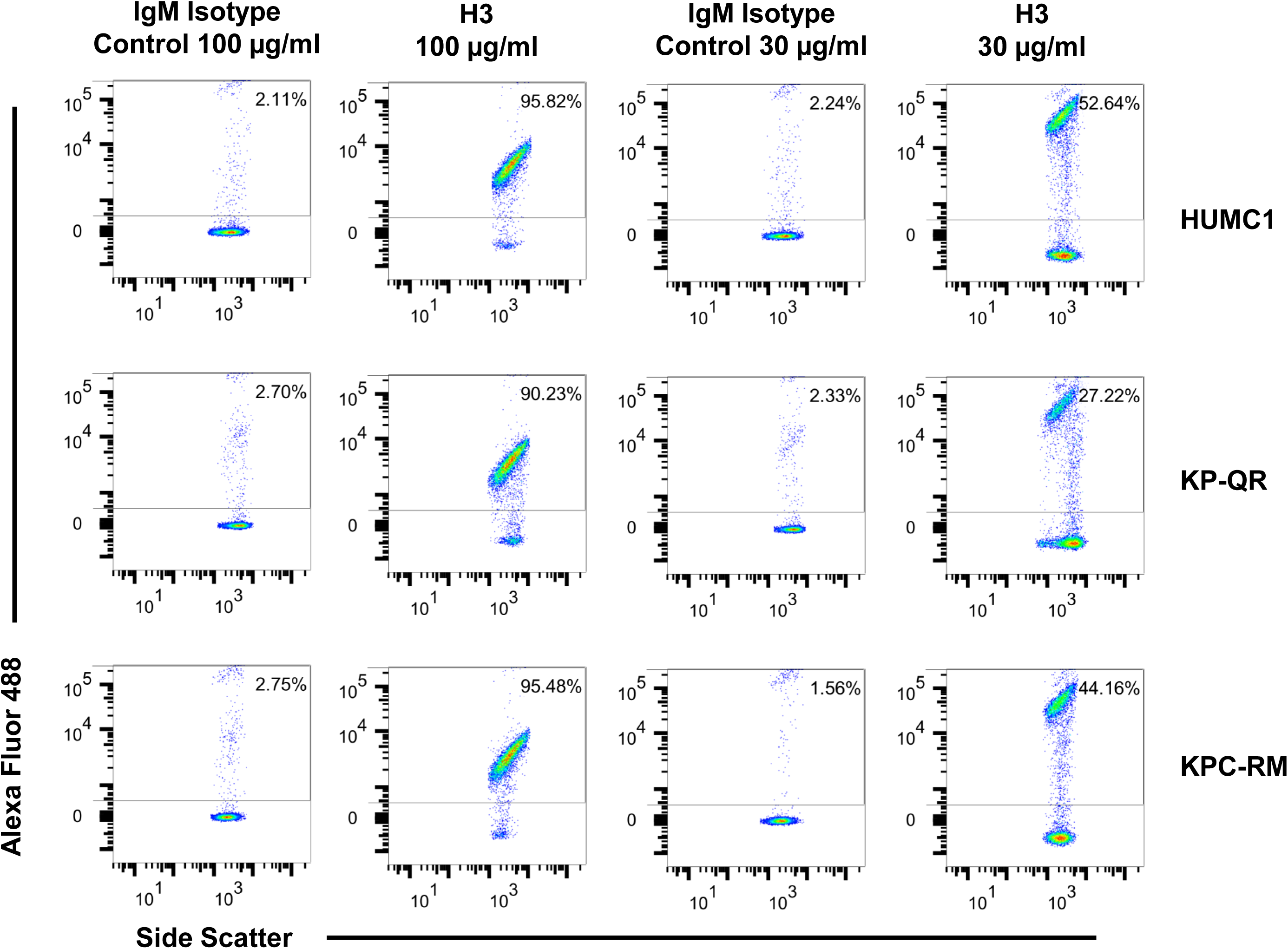
mAb clone H3 targeting Hyr1p peptide 5 binds GNB in a dose dependent manner. mAb clones (and IgM isotype controls) were evaluated for binding to HUMC-1, KP-QR, and KP-RM, and the extent of binding quantified by flow cytometry after staining the antibodies with alexa 488-conjugated secondary antibody. Data was represented in a scatter-plot highlighting the percentage of bacteria that were bound by the Abs.

### mAbs protect host cells from damage by GNB

Our previous studies demonstrated that anti-Hyr1 pAbs could not only bind to, but also inhibit *A. baumannii*’s ability to damage mammalian cells ^14^. Thus, we hypothesized that binding of GNB with mAbs would similarly block the invasion and damage of host cells by the MDR GNB, *A. baumannii* (HUMC1), KP-QR and KPC-RM. Indeed, we found that mAb clones H3 and H4 at 30 μg/ml reduced the ability of HUMC1 and KP-QR to damage lung A549 cells by ∼70-90%, in the presence of 15-30 μg/ml of H3 and H4. The carbapenamase resistant KP, KPC-RM was slightly more resistant to the mAbs, displaying reduction in damage of 40% in 15 μg/ml of the mAbs; the higher 30 μg/ml of mAbs prevented 70% of damage to A549 cells by the KPC (**Fig 3A**). The two mAb also protected the lung cell line from damage by other clinical isolates of GNB such as, HUMC-6 and KPC-8 (**S2**). Consistent with these results, both mAb at 15 *µ*g/ml resulted in ∼40-70% inhibition of *A. baumannii* HUMC1- or KP-QR-mediated damage to the primary human vascular endothelial cells (HUVEC), respectively (**Fig 3B**). However, it took a higher mAb concentration, 30 *µ*g/ml, to protect HUVECs from KPC-RM. The mAbs were also found to be active in protecting HUVEC cells from two other clinical isolates, *A. baumannii* HUMC-6 and KPC-8 (**Fig S2**). Furthermore, we found that mAb H3 was capable of directly killing KP-QR and HUMC1, but not another drug resistant GNB *Pseudomonas aeruginosa* (PA) which does not have surface proteins homologous to Hyr1p (**Fig 3C**). Interestingly, KPC-RM was completely resistant to direct killing by the both the mAb clones H3 and H4 (**S3A**). Overall, these results show that mAbs raised against Hyr1p peptide 5 bind to MDR *A. baumannii* and *K. pneumoniae* strains, block their ability to damage host cells, and directly kill the bacteria, *in vitro*.

**Figure 3.**
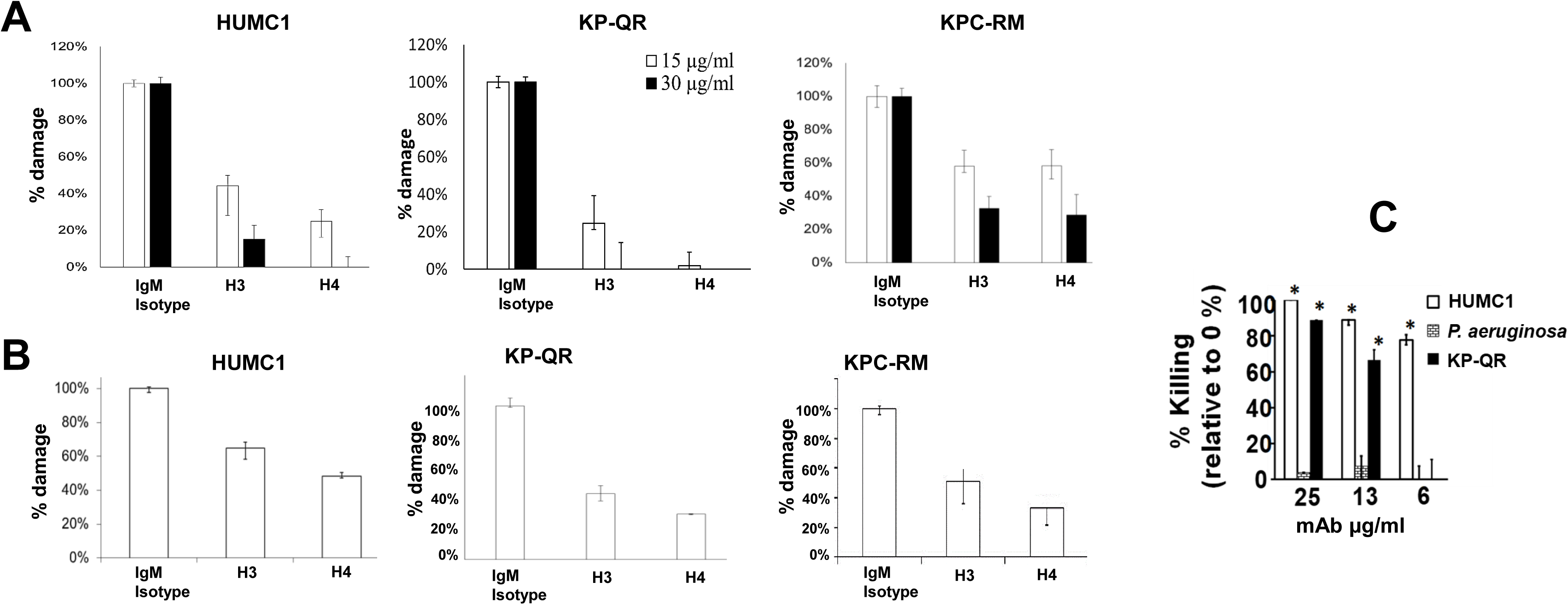
mAb targeting Hyr1p peptide 5 prevents HUMC1, KP-QR and KP-RM-induced alveolar epithelial (A549) cell damage and primary human endothelial cell damage (HUVEC), and directly kill bacteria. HUMC1, KP-QR and KPC-RM-induced A549 cell injury in the presence of 15 or 30 μg/ml of an isotype matching IgM, mAb H3, or mAb H4 **(A)**. Damage to HUVEC cells by HUMC1 and KP-QR (in presence of15 μg/ml of H3 and H4), also KPC-RM (30 μg/ml of the mAbs) **(B)**. Cell injury was determined by ^51^Cr-release assay^15^after 48 or 24 h for HUMC1 and KP, respectively. *\*P* <0.001 vs. Control IgM. n=12 from 3 experiments. Effect of mAb on the viability of HUMC1, PA or KP-QR *in vitro* **(C)**. Bacterial cells (1 × 10^5^ cells) in MHII medium were incubated in 96-well plate at 37°C for 20 h with varying concentrations of mAb H3. Killing activity of H3 was enumerated by CFU (n= 3 per arm) expressed as % killing relative to cells without H3. \**P*<0.005 vs. no H3 by Wilcoxon Rank Sum test.

### Anti-Hyr1 mAbs protect mice from pulmonary infection caused by *K. pneumoniae* and *A. baumannii*

Given that the mAbs bound to GNB and neutralized their ability to damage host cells *in vitro*, we tested their ability in protecting mice from GNB infections. Pneumonia is a major manifestation of the disease caused by both *K. pneumoniae* and *A. baumannii* ^*15-18*^. Thus, we evaluated H3 and H4 for their potential to protect against pulmonary disease caused by *A. baumannii* HUMC1. Although benign in immunocompetent individuals, *A. baumannii* can cause life-threatening pneumonia in immunosuppressed hospitalized patients ^18^. Thus, we utilized a neutropenic mouse model to infect mice with HUMC1, via inhalation. The mAb were administered on Day +1, and +4, relative to infection mice at a dose of 30 μg administered intraperitoneally. Placebo mice were treated similarly with an isotype matched control IgM. Compared to the placebo-treated mice, mAb H4 resulted in a high 70% overall survival versus 20% overall survival for placebo mice. Impressively, complete protection (100% survival) was elicited in mice receiving H3 mAb (**Fig 4A**). Surviving mice appeared healthy on Day +21 when the experiment was terminated.

**Figure 4.**
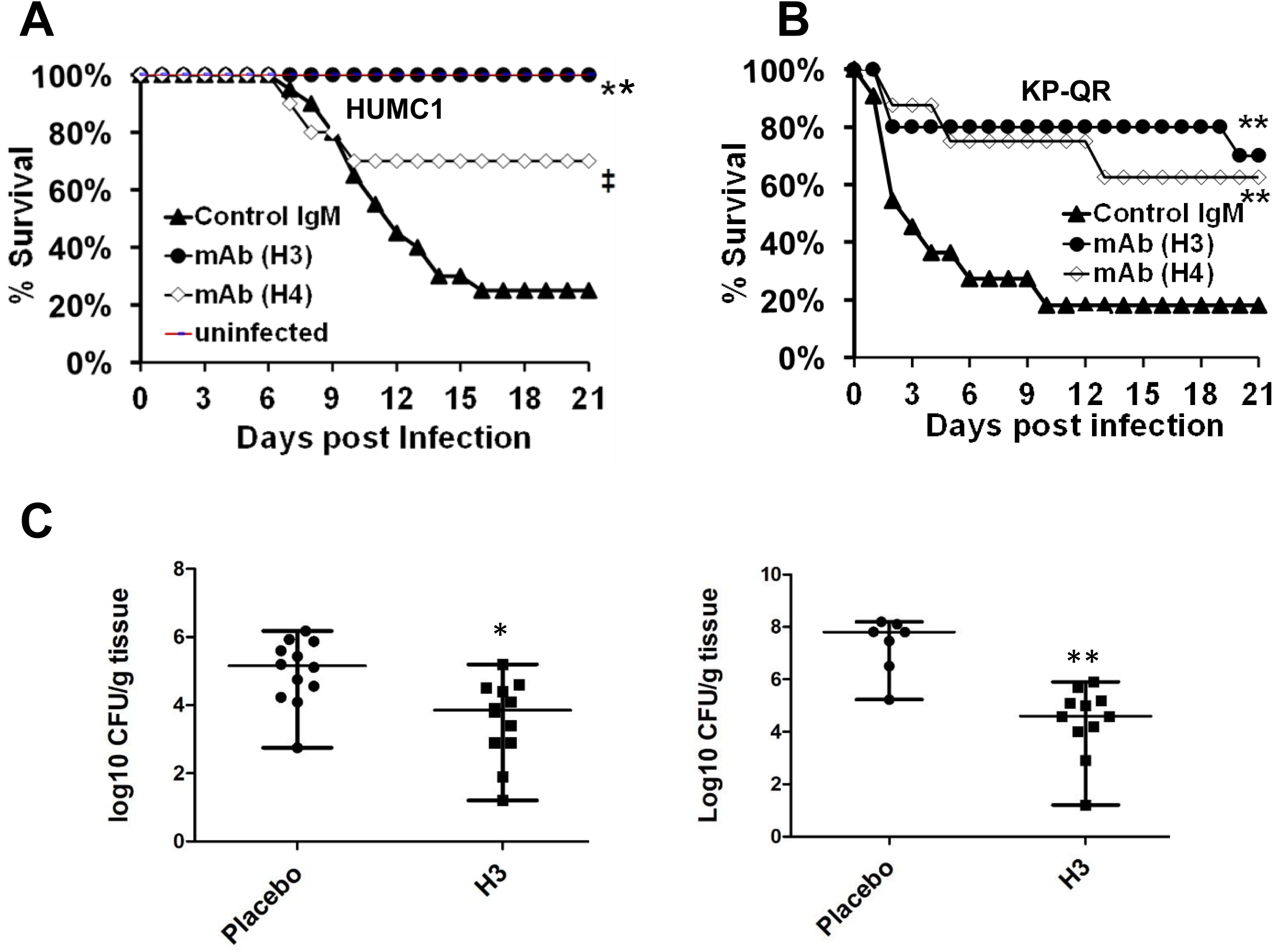
mAb clones H3 and H4, targeting Hyr1p peptide 5, protect against HUMC1 and KP-QR induced pneumonia. Immunosuppressed CD-1 male mice (n=10/group) were infected with HUMC1 via inhalation (**A**). Immunocompetent mice (n=10/group from 2 experiments) were infected intratracheally with KP-QR (ave. 3.4 × 10^7^) (**B**). I.P. treatment with mAb H3, H4, or isotype-matched control IgM started 24 h and repeated at 96 h post infection (30 μg each dose). *P<0.01, ‡P=0.06, and **P<0.001 vs. control IgM by Log rank test. For CFU measurement, mAb were administered 6 h and 3 day post infection, and lungs harvested from mice at day +4 (for HUMC1) and day +2 (for KP-QR) post infection (**C**).

We next evaluated the efficacy of mAb in a similar pulmonary mouse model of *K. pneumoniae* infection. Our *in vivo* optimization studies have shown that KP-QR exhibits pronounced lethality even in healthy non-immunosuppressed mice, while KPC-RM are avirulent despite high cell numbers used for infection (**S3B**). Thus, we evaluated the protective effect of mAb against the KP-QR-mediated pneumonia in mice. Immunocompetent mice were infected intratracheally with KP-QR, and therapeutically treated twice (1 day and 4 days after infection) with mAbs, or isotype matching control IgM. Almost 60% of the mice treated with H4 survived the pulmonary infection by KP-QR (p<0.06), while yet again, H3 showed a significantly improved protection (trending to 100% survival in mice) (**Fig 4B**). Surviving mice appeared healthy at day 21 when the experiment was terminated.

At different time points during infection (+2 days for KP-QR and +4 days for HUMC1), lungs from H3 vaccinated mice were harvested for measurement of bacterial burden. Corroborating the survival data, in comparison to treatment with isotype matching IgM controls, H3 resulted in 1.5- and 3-log reduction in lung bacterial burden of HUMC1 and KP-QR, respectively (P<0.01) (**Fig 4C**).

Together, these results demonstrate that therapeutic mAbs exhibit translational potential as a novel treatment option for GNB-associated pneumonia in different hosts (immunocompetent as well as neutropenic).

## Discussion

Different pathogens inhabit common settings in the host and depend on similar strategies for adaptation to stress, to proliferate, and to cause pathogenesis. Indeed the fungus *C. albicans* and certain GNB, such as *A. baumannii* and *K. pneumoniae* infect the same patient populations, encompassing those who are immunocompromised and hospitalized, or patients suffering from burn and surgical wounds ^7,19,20^. In fact, *Candida* species airway colonization together with *A. baumannii*, during ventilator-associated pneumonia (VAP), are common among ICU patients, and identified as an independent risk factor for development of *A. baumannii* VAP ^21^. Similarly, *Candida* and *Klebsiella* are the most frequent pathogens of the respiratory tract of patients with chronic obstructive pulmonary disease (COPD) ^22,23^. Above and beyond their association with each other, *Candida* and GNB also individually cause healthcare-associated infections, often leading to life threatening diseases. The GNB, in particular, have evolved over the years into multidrug resistant entities causing infections that are often incurable ^24^. In the last two decades, the incidence of MDR GNB including *A. baumannii* and *K. pneumoniae* has been so high, that the standard of public health in many parts of the world is considered equivalent to the pre-antibiotic era ^24^. Hence, newer approaches that go further than simply the discovery of new antibiotics are needed to combat the crisis of drug resistance.

Our group uses advanced computational molecular modeling and bioinformatic approaches to discover novel vaccine antigen candidates that target more than one high priority human pathogens ^14,25,26^. This strategy, known as unnatural or heterologous immunity, has been previously applied in the development of viral and bacterial vaccines in which an antigen protects against another pathogen from the same or from a different kingdom ^25^. We have previously validated this approach by demonstrating cross kingdom immuno-protection against *C. albicans* and *S. aureus*, in which the *C. albicans* cell surface adhesin/invasion proteins (Als family of proteins) share structural and functional homology with MSCRAMMs of *S. aureus* (e.g. clumping factor A) ^27^. A recombinant form of N-terminus of the Als3p (rAls3p-N) elicits robust T- and B-cell responses and protects mice from both *Candida* and MRSA infections ^26,28-33^. Most recently, we reported that another hyphal cell surface protein of *C. albicans*, Hyr1p, has 3-D structural homologies with candidate antigens of the MDR GNB *A. baumannii* ^*14*^. Indeed, using different mouse models, active or passive immunization (with pAb) targeting either Als3p or Hyr1p, protected mice from *S. aureus* or *A. baumannii* infections, respectively ^14,26,28^. In particular, antibodies against one specific surface exposed and highly antigenic15-mer peptide of Hyr1p offered the highest protection to host cells from *A. baumannii* both *in vitro* and *in vivo* ^14^.

Encouraged by the potential of the pAb, and to further the clinical scope of our studies, we produced and developed mAbs against this highly antigenic peptide of Hyr1. Similar to pAbs, this study showed that mAbs blocked the pathogenesis of both *A. baumannii-* and *K. pneumoniae*-mediated host cell damage and protected mice from pulmonary infections caused by MDR *A. baumannii* or *K. pneumoniae*. Initial functional assays revealed that four different clones of mAbs (H1-H4) recognized the two genera of bacteria in *in vitro* binding assays, with high sensitivity. This binding capacity was extended also to include several MDR clinical isolates of *A. baumannii* and *K. pneumoniae*.

The ability of the mAbs (H3 and H4) to block GNB-mediated damage of host cells, was more pronounced in *A. baumannii* HUMC1, *A. baumannii* HUMC6, and *K. pneumoniae* KP-QR, rather than KPC-RM or KPC-8. Four factors, capsule, lipopolysaccharide, fimbriae, and siderophores, have been identified as important for pathogenesis, and resistance of hypervirulent KP strains to antibiotics ^34^. Thus, resistance of KPC to killing by mAb could be due to the difference in the capsule structure, or their reported ability to produce a larger repertoire of siderophores ^34^. Indeed, our recent report does emphasize the importance of anti-Hyr1p peptide 5 polyclonal antibodies in blocking iron uptake, leading to killing of the GNB *A. baumannii* ^14^.

Nevertheless, almost a 70% protection from cellular damage was provided by the mAbs in all GNB tested, making mAb a potential therapeutic agent with capacity to block virulence of GNB. This point becomes even more important considering the mAb (just like the pAb ^14^) can directly kill *Acinetobacter* and *Klebsiella*, but not another GNB *Pseudomonas* whose proteins were not identified to be homologous to Hyr1p ^14^. Our recent report on bioinformatic, homology and energy-based modeling strategies described that *C. albicans* Hyr1p shares striking similarity to *A. baumannii* FhaB protein, and anti-peptide #5 pAb could bind to FhaB as well as two other proteins on *A. baumannii* on 2 dimensional Western blotting assays ^14^. The other two proteins included the outer membrane protein OmpA, and a ferric siderophore outer membrane binding protein (TonB)^14^. Not surprisingly, the three proteins are conserved in *K. pneumoniae* displaying >60% homologies to their *Acinetobacter* counterparts (protein sequence NCBI blast alignment) (**S4**). Whether these proteins have a significant role in virulence or nutrient uptake – and hence blocking their function would contribute the killing mechanisms afforded by the mAbs, is the subject of ongoing research by our group.

Because the mAbs significantly blocked the capacity of GNB to damage host cells, we further evaluated their potential to protect against pulmonary infections caused by the two respective bacteria. H4 protected >60% of mice from succumbing to pulmonary infection by KP-QR compared to placebo that only had a 20% survival rate. In fact, the mAb H3 provided total (100%) protection to mice from *A. baumannii* HUMC1 infection, similar to that conferred by pAb^14^. This indicates that mAb likely recognize specific targets on the bacterium that when neutralized, attenuate its ability to survive in the host and cause disease. Certainly, therapeutic treatment by mAbs interfered significantly with the dissemination of KP-QR and HUMC1 to their target organs within the first 2 to 4 days of treatment, versus those treated with control antibodies. This is an evidence for the robustness of the antibodies in abrogating pathogenesis, early into the onset of infection.

An advantage of using mAbs in infectious diseases is their well-documented long half-life which can be in excess of 21 days ^35,36^. Since the patient population at risk of developing infections with *Acinetobacter, Klebsiella*, and *Candida* are well-defined, the property of mAbs long half-life can prove to be critical in reducing the use of antibiotics and hence development of drug resistance. For example, mAbs can be used to prophylax patients at risk of MDR GNB. Another envisioned usage of these mAbs is their administration as adjunctive therapy with antibiotics. In this respect, we have demonstrated synergy of anti-peptide 5 pAb with imipenem or with colistin in killing *A. baumannii* at reduced minimum inhibitory concentrations (MIC) of both antibiotics^14^. Similarly, the anti-peptide 5 pAb synergistically acted with colistin in abrogating biofilm growth of *A. baumannii*^14^.

In summary, we have demonstrated that mAbs raised against anti-peptide 5 of Hyr1p, specifically recognize *A. baumannii* and *K. pneumoniae*, and disrupt their ability to cause damage to host cells. Importantly, these mAbs protect mice from lethal pulmonary infections by these two GNB, and have the potential to be used as prophylactic or adjunctive therapy to improve the standard of care for immunosuppressed patients susceptible to MDR *A. baumannii* or *K. pneumoniae*.

## Material and methods

### Bacterial strains and growth conditions

The bacterial strains used in this study are clinical isolates collected from Harbor-UCLA Medical Center (Torrance, CA). *A. baumannii* strains; HUMC-1, HUMC-6 were separated from patients’ sputum, and HUMC-12 separated from patients’ wound are extensively drug resistant to all antibiotics, except colistin and tigecycline. *K. pneumoniae* strains were categorized into KPC or non-KPC isolates. The KPC-RM, KPC-6, KPC-7, KPC-8 isolates resistant to carbapenem antibiotics, possess *bla* KPC plasmid gene and separated from patients’ sputum. While KP-QR; is a non-KPC multi-drug resistant isolate separated from patients’ sputum and resistant to Gentamicin, Kanamycin and Ampicillin/sulbactam antibiotics. All bacteria were cultured in Tryptic soya broth (TSB) overnight at 37°C with shaking at 200 rpm. To obtain a log-phase bacterial suspension; overnight cultured bacteria were passaged in a fresh media (1:100) at 37°C with shaking for 3 hours or until the cell concentration reached and OD_600_ of 0.5 (∼ 2 × 10^8^ cells/ml) for both *K. pneumoniae* and *A. baumannii* isolates. The bacteria were diluted to the desired concentration from this stock.

### Generation of mAbs

30 *µ*g of rHyr1 peptide #5 (commercially purchased) in 1 mg/ml Alum was used to immunize Balb/c mice (n=10). The mice were boosted two times every 2 weeks with the same antigen concentration. Two weeks after the last boost; antibody titer was determined by ELISA. The spleens were collected and the splenocytes were fused with Hypoxanthine-guanine phosphoribosyltransferase (HGPRT) negative murine myeloma cells at ratio 5:1 by slowly adding poly ethylene glycol (PEG) to the cells pellet and Protein Free Hybridoma Media (PFHM) (Gibco, 12040077) supplemented with 20% Heat inactivated Fetal Bovine Serum (FBS) (Corning, 35-016-CV) was added. The cells were spun and re-suspended in 20% FBS PFHM, and were incubated in 24-well plate at 37°C with 5% CO_2_ for 48 hours. The media was replaced with Hypoxanthine-Aminopterin-Thymidine (HAT) selection media for 8 days, then 20% FBS PFHM HT media for 2 weeks. The grown hybridoma clones were diluted by microdilution in microtiter plates to achive one cell per well and kept culturing in 10 % FBS PFHM. The supernatant from the grown clones were tested for anti-Hyr1 antibodies using ELISA. The selected positive and stable clones were cultured in PFHM without FBS and the cell numbers were adjusted to be 2 × 10^5^/ml for optimum production of mAbs.

### Characterization and Purification of mAbs

The supernatant containing the antibodies were concentrated using 100 KDa cutoff centrifugal concentrating tube (Amicon, UFC910024). HiTrap HP column (GE Healthcare, 17511001) were used to purify the concentrated mAbs, and then buffer exchanged with endotoxin-free Dulbecco PBS without Calcium or magnesium (Gibco, 14190250). Endotoxin was tested using a chromogenic limulus amebocyte lysate assay (BioWhittaker Inc.). Finally the mAbs isolated on an SDS-PAGE gel.

### Surface staining and binding assay

Bacterial cells (5 ×10^6^ cells) re-suspended in 2% FBS-PBS were incubated with anti-Hyr1 mAbs, pAbs or isotype matching control for 2 hours at higher conc. (100*µ*g/ml) or lower conc. (30*µ*g and 3 *µ*g/ml). The bacterial cells were washed three times with cold 2% FBS-PBS. The bound anti-Hyr1 pAbs and mAbs were detected with anti-mouse or anti-rabbit FITC-labelled secondary antibodies. The unbound antibodies were washed out with cold 2% FBS-PBS before measuring the fluorescent stained bacterial cells using flow cytometry.

### Cell damage assay

The in vitro ability of mAbs to protect either A549 cells or HUVEC from damage caused by direct contact with bacteria was measured using ^51^Cr release assay, modified from previous method^37^. Isolation of HUVEC cells were performed in the lab under a protocol approved by institutional IRB. Because umbilical cords are collected without donor identifiers, the IRB considers them medical waste not subject to informed consent.

Lung A549 cells and HUVEC cells were incubated overnight in 24-well plates with F-12K or RPMI medium containing 6 μCi/well of Na_2_ ^51^CrO_4_ (ICN Biomedicals, Irvine, CA). The next day, unincorporated tracer was aspirated and the wells were rinsed three times with warm HBSS. One milliliter of HBSS containing HUMC1 or KP-QR was added to each well at an MOI of 1:100 (cells to bacteria), and the plate was incubated for 3 h at 37°C in 5% CO_2_. At the end of the incubation, 0.5 ml of medium was gently aspirated from each well, after which the endothelial cells were lysed by the addition of 0.5 ml of 6 N NaOH. The lysed cells were aspirated, and the wells were rinsed twice with RadioWash (Atomic Products, Inc., Shirley, N.Y.). These rinses were added to the lysed cells, and the ^51^Cr activity of the medium and the cell lysates was determined. Control wells containing HBSS but no organisms were processed in parallel to measure the spontaneous release of 51Cr. After corrections were made for the differences in the incorporation of 51Cr in each well, the specific release of 51Cr was calculated by the following formula: (2 X experimental release - 2 X spontaneous release)/ (total incorporation - 2 X spontaneous release).

### Animal infection models

All procedures involving mice were approved by the IACUC of the Los Angeles Biomedical Research Institute at Harbor-UCLA Medical Center, according to the NIH guidelines for animal housing and care. 4-6 week aged CD-1 immunocompetent male mice were used for *Klebsiella* (KP-QR) intratracheal infection; immunosuppressed mice infected with *A. baumannii* (HUMC-1) by aerosolization chamber to induce pneumonia by inhalation. Mice were immunosuppressed by administrating cyclophosphamide [200 mg/kg] (i.p.) and cortisone acetate [250 mg/kg] (s.c.) on days −2, +3, and +8 relative to infection as previously described^37^. A total of 30 *µ*g of mAbs or isotype matching control were administrated (i.p.) on day (+1) and on day (+4) post infection, and survival of mice served as an endpoint. For quantitative measurement of bacterial burden, mAb were administered 6 hours post infection and boosted on day (+3). Mice were euthanized on day (+2) (for *A. baumannii*) and on day (+4) (for *K. pneumoniae*), lungs harvested aseptically, homogenized, and the bacterial burden determined by quantitative culturing.

### Statistical analysis

Percentage of cell damage and tissue bacterial burden were compared using non-parametric Man Whitney test. Log Rank test was used to determine the difference in survival studies. P values of <0.05 considered significant.

---

**Supplemental figure 1.**
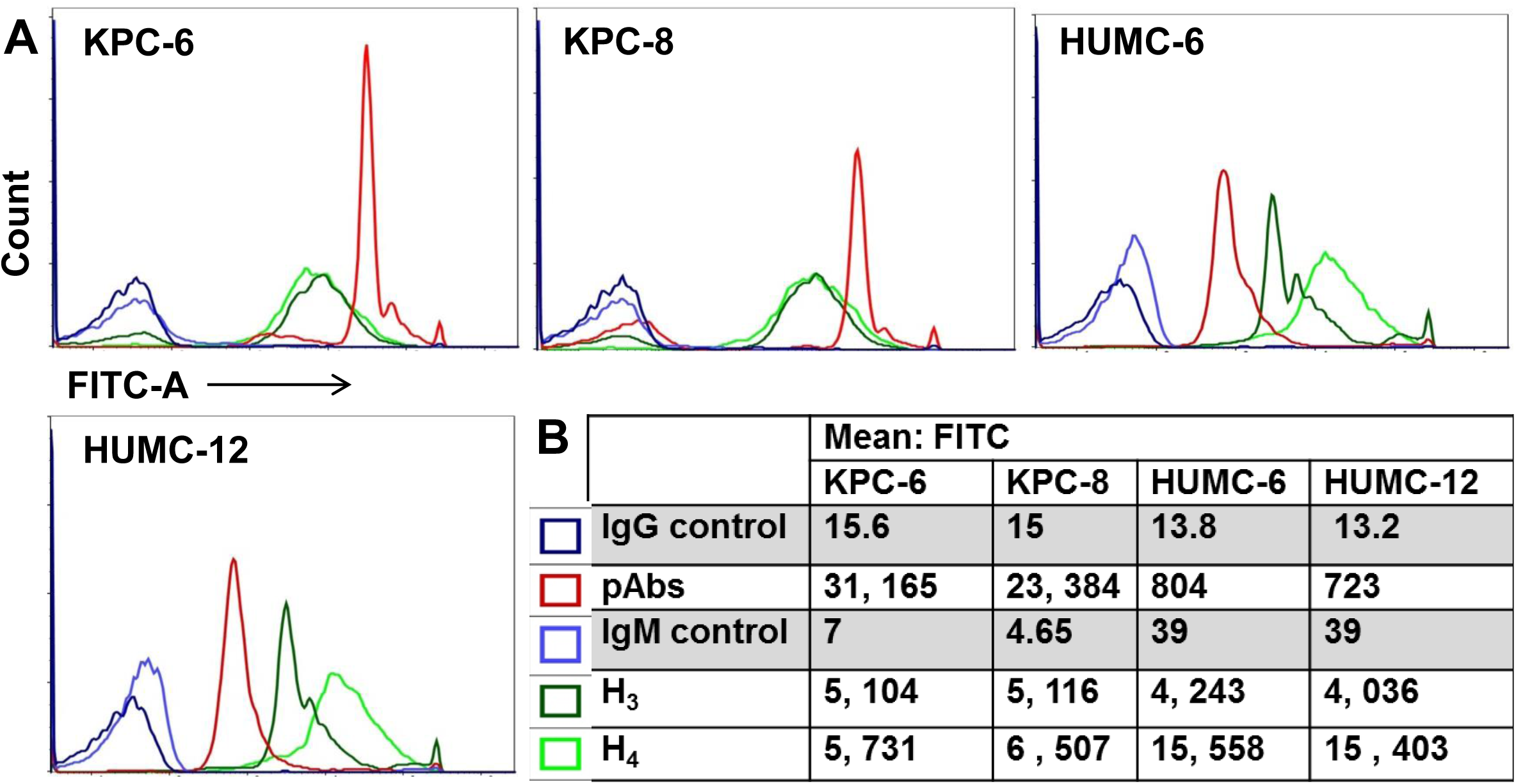
Binding of the mAb clones H3 and H4, targeting Hyr1p peptide 5, to clinical isolates of GNB: mAb clones H3 and H4 (and polyclonal antibodies, as well as isotype-matched control IgM) were evaluated for binding to KPC-6, KPC-8, HUMC-6 *and* HUMC-12 at a concentration of at 100 *µ*g/ml. The extent of binding was quantified by flow cytometry after staining the bound antibodies with alexa 488-conjugated secondary antibody. The degree of binding was visualized by a shift in the peaks in the anti-Hyr1 IgM binding conditions versus the control antibodies (A). Data was also quantified and shown as mean fluorescence intensity of the Ab-bound bacteria (**B**).

**Supplemental figure 2.**
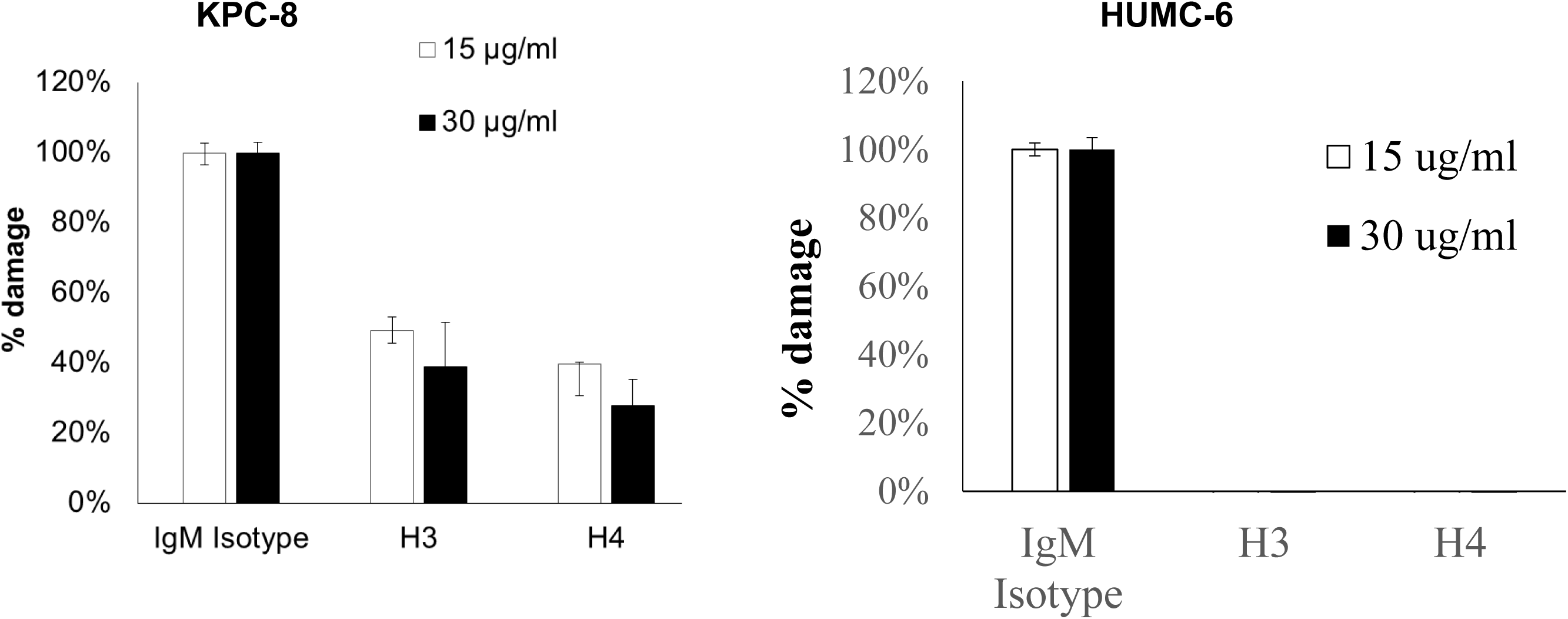
mAb clones H3 and H4 protect lung A549 cells from damage by GNB. KPC8 and HUMC-6 were incubated individually with Mab H3 and H4 for an hour, prior to challenging ^51^Cr-labeled lung A549 cells for 20 h. Extent of damage was assessed by quantifying release of ^51^Cr from the mammalian cells. % damage was normalized to isotype matching control after subtracting spontaneous cell damage. N= 12 per each group from 3 independent experiment p<0.0001 vs. IgM isotype-matched controls.

**Supplemental figure 3.**
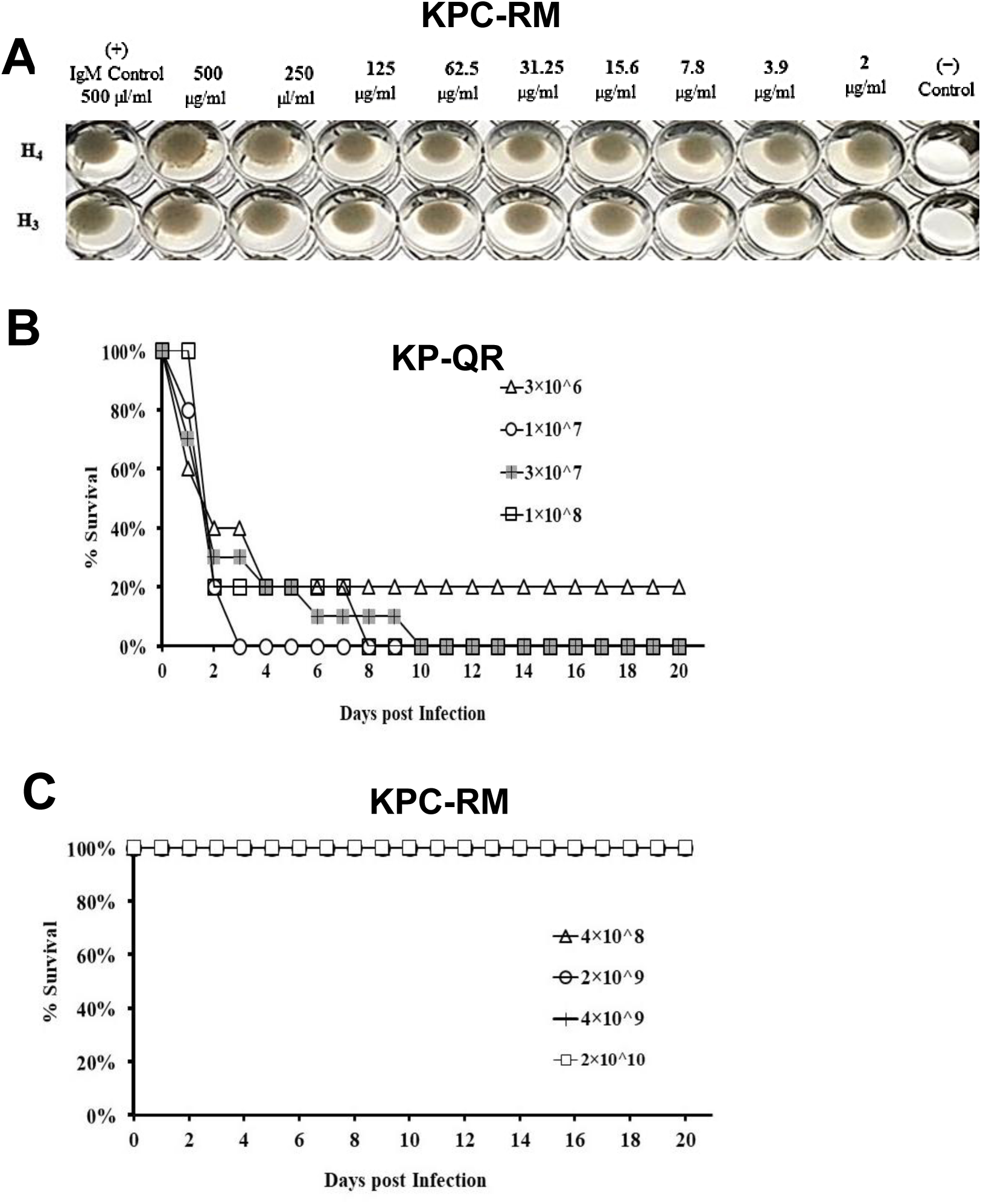
mAb-mediated killing of KP strains *in vitro*, and survival curve of mice infected separately with different inocula of KP-QR and KPC-RM isolates. mAb clones H1, H2, H3 and H4 were serially diluted two-fold in Mueller Hinton broth, in a 96 well plate. The concentration of mAb tested ranged from 500 *µ*g/ml to 2 *µ*g/ml. Isotype-matched IgM at 500 *µ*g/ml served as a control isotype-matching control. Killing of the mAb were tested against 10^5^ cells/ml of the respective KP strains, after a 24 h incubation at 37°C, and were subject to visual inspection (**A**). Immunocompetent mice were infected via intratracheal route. N=5 per each group, and survival was chosen as the primary end point (**B**).

**Supplemental figure 4.**
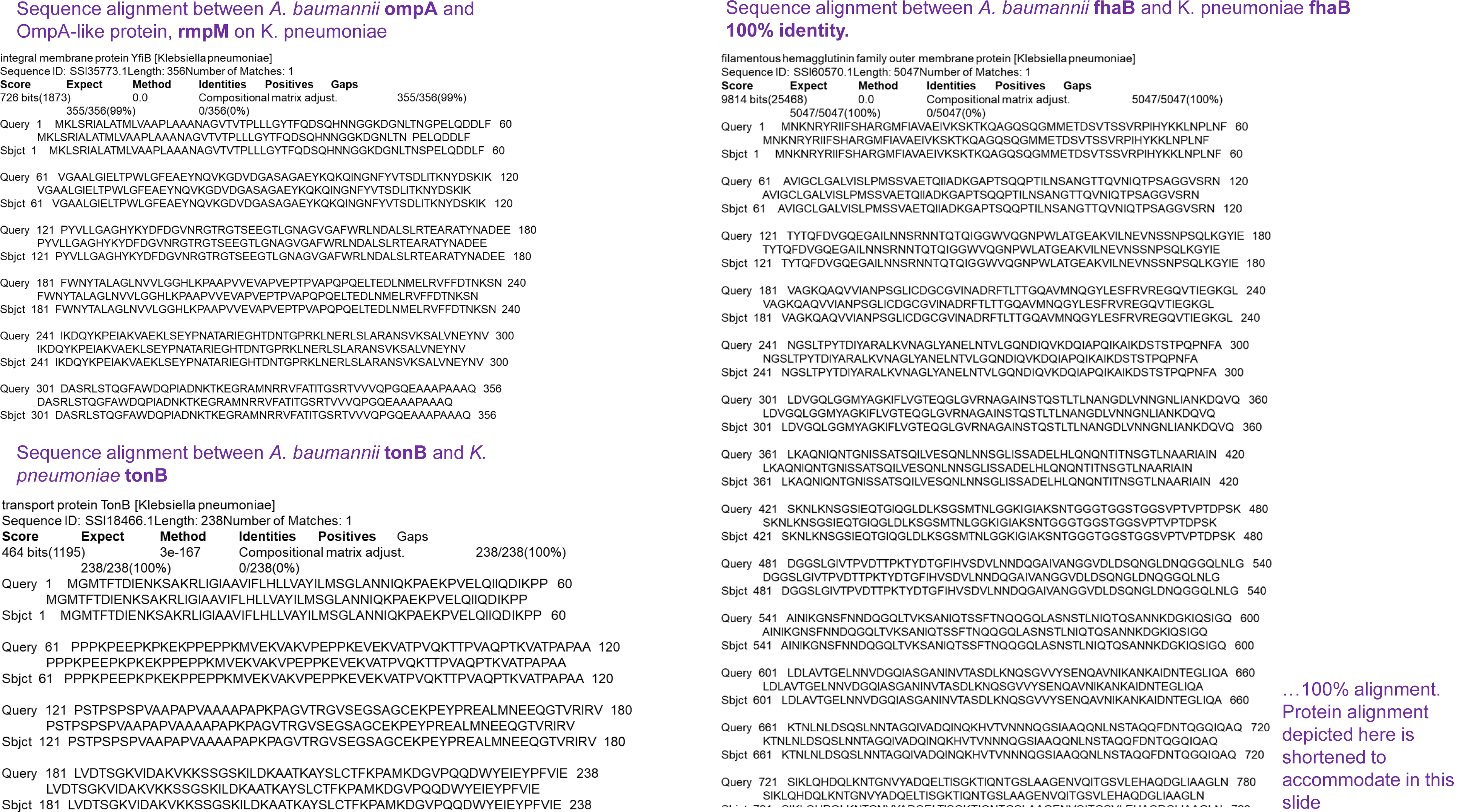
Sequence alignments of OmpA, TonB and FhaB between AB and KP. The three mentioned proteins of AB had 99-100% identity with the respective proteins of KP.

## References

1 Perez, F. et al. Global challenge of multidrug-resistant Acinetobacter baumannii. Antimicrob Agents Chemother 51, 3471–3484, DOI: AAC.01464-06 [pii]10.1128/AAC.01464-06 (2007).

2 Falagas, M. E., Karveli, E. A., Siempos, I. & Vardakas, K. Z. Acinetobacter infections: a growing threat for critically ill patients. Epidemiol Infect 136, 1009–1019, DOI: S0950268807009478 [pii]10.1017/S0950268807009478 (2008).

3 Karageorgopoulos, D. E. & Falagas, M. E. Current control and treatment of multidrug-resistant Acinetobacter baumannii infections. Lancet Infect Dis 8, 751–762, DOI:S1473-3099(08)70279-2 [pii]10.1016/S1473-3099(08)70279-2 (2008).

4 Higgins, P. G., Dammhayn, C., Hackel, M. & Seifert, H. Global spread of carbapenem-resistant Acinetobacter baumannii. J Antimicrob Chemother 65, 233–238, DOI:dkp428 [pii]10.1093/jac/dkp428.

5 DOI, Y., Husain, S., Potoski, B. A., McCurry, K. R. & Paterson, D. L. Extensively drugresistant Acinetobacter baumannii. Emerg Infect Dis 15, 980–982 (2009).

6 Hoffmann, M. S., Eber, M. R. & Laxminarayan, R. Increasing resistance of acinetobacter species to imipenem in United States hospitals, 1999-2006. Infect Control Hosp Epidemiol 31, 196–197, DOI:10.1086/650379.

7 Rosenthal, V. D. et al. International Nosocomial Infection Control Consortium (INICC) report, data summary for 2003-2008, issued June 2009. Am J Infect Control 38, 95–104 e102, DOI:S0196-6553(09)00952-3 [pii] 10.1016/j.ajic.2009.12.004.

8 Lautenbach, E. et al. Epidemiology and impact of imipenem resistance in Acinetobacter baumannii. Infect Control Hosp Epidemiol 30, 1186–1192, DOI:10.1086/648450 (2009).

9 Mammina, C. et al. Ongoing spread of colistin-resistant Klebsiella pneumoniae in different wards of an acute general hospital, Italy, June to December 2011. Euro Surveill 17 (2012).

10 Brink, A. J. et al. Emergence of OXA-48 and OXA-181 carbapenemases among Enterobacteriaceae in South Africa and evidence of in vivo selection of colistin resistance as a consequence of selective decontamination of the gastrointestinal tract. J Clin Microbiol 51, 369–372, DOI:10.1128/JCM.02234-12 (2013).

11 Giordano, C. et al. Expansion of KPC-producing klebsiella pneumoniae with various mgrB-mutations giving rise to colistin-resistance: the role of ISL3 on plasmids. International Journal of Antimicrobial Agents, DOI:https://DOI.org/10.1016/j.ijantimicag.2017.10.011 (2017).

12 Luo, G. et al. Candida albicans Hyr1p confers resistance to neutrophil killing and is a potential vaccine target. J Infect Dis 201, 1718–1728, DOI:10.1086/652407 (2010).

13 Luo, G., Ibrahim, A. S., French, S. W., Edwards Jr., J. E. & Fu, Y. Active and Passive Immunization with rHyr1p-N Protects Mice against Hematogenously Disseminated Candidiasis. PLoS One 6, e25909, DOI:DOI:10.1371/journal.pone.0025909 (2011).

14 Uppuluri, P. et al. The Hyr1 protein from the fungus Candida albicans is a cross kingdom immunotherapeutic target for Acinetobacter bacterial infection. PLoS Pathog 14, e1007056, DOI:10.1371/journal.ppat.1007056 (2018).

15 Trouillet, J. L. et al. Ventilator-associated pneumonia caused by potentially drug-resistant bacteria. Am J Respir Crit Care Med 157, 531–539 (1998).

16 Park, D. R. The microbiology of ventilator-associated pneumonia. Respir Care 50, 742-763; discussion 763-745 (2005).

17 Guidelines for the management of adults with hospital-acquired, ventilator-associated, and healthcare-associated pneumonia. Am J Respir Crit Care Med 171, 388–416, DOI:171/4/388 [pii] 10.1164/rccm.200405-644ST (2005).

18 Caricato, A. et al. Risk factors and outcome of Acinetobacter baumanii infection in severe trauma patients. Intensive Care Med 35, 1964–1969, DOI:10.1007/s00134-009-1582-5 (2009).

19 Peleg, A. Y., Seifert, H. & Paterson, D. L. Acinetobacter baumannii: emergence of a successful pathogen. Clin Microbiol Rev 21, 538–582, DOI:21/3/538 [pii] 10.1128/CMR.00058-07 (2008).

20 Gaddy, J. A., Tomaras, A. P. & Actis, L. A. The Acinetobacter baumannii 19606 OmpA protein plays a role in biofilm formation on abiotic surfaces and in the interaction of this pathogen with eukaryotic cells. Infect Immun 77, 3150–3160, DOI:IAI.00096-09 [pii] 10.1128/IAI.00096-09 (2009).

21 Tan, X. et al. Candida spp. airway colonization: A potential risk factor for Acinetobacter baumannii ventilator-associated pneumonia. Med Mycol 54, 557–566, DOI:10.1093/mmy/myw009 (2016).

22 Przybylowska, D. et al. Evaluation of Genetic Diversity of Candida spp. and Klebsiella spp. Isolated from the Denture Plaque of COPD Patients. Adv Exp Med Biol 955, 1–8, DOI:10.1007/5584_2016_68 (2017).

23 Su, J. et al. Sputum Bacterial and Fungal Dynamics during Exacerbations of Severe COPD. PLoS One 10, e0130736, DOI:10.1371/journal.pone.0130736 (2015).

24 Ventola, C. L. The antibiotic resistance crisis: part 1: causes and threats. P T 40, 277–283 (2015).

25 Yeaman, M. R. et al. Applying Convergent Immunity to Innovative Vaccines Targeting Staphylococcus aureus. Front Immunol 5, 463, DOI:10.3389/fimmu.2014.00463 (2014).

26 Yeaman, M. R. et al. Mechanisms of NDV-3 vaccine efficacy in MRSA skin versus invasive infection. Proc Natl Acad Sci U S A 111, E5555–5563, DOI:10.1073/pnas.1415610111 (2014).

27 Sheppard, D. C. et al. Functional and structural diversity in the Als protein family of Candida albicans. J Biol Chem 279, 30480–30489 (2004).

28 Lin, L. et al. Th1-Th17 cells mediate protective adaptive immunity against Staphylococcus aureus and Candida albicans infection in mice. PLoS Pathog 5, e1000703, DOI:10.1371/journal.ppat.1000703 (2009).

29 Spellberg, B. et al. The antifungal vaccine derived from the recombinant N terminus of Als3p protects mice against the bacterium Staphylococcus aureus. Infect Immun 76, 4574–4580, DOI:IAI.00700-08 [pii] 10.1128/IAI.00700-08 (2008).

30 Lin, L. et al. Immunological surrogate marker of rAls3p-N vaccine-induced protection against Staphylococcus aureus. FEMS Immunol Med Microbiol, DOI:FIM531 [pii] 10.1111/j.1574-695X.2008.00531.x (2009).

31 Spellberg, B. et al. Antibody titer threshold predicts anti-candidal vaccine efficacy even though the mechanism of protection is induction of cell-mediated immunity. J Infect Dis 197, 967–971, DOI:10.1086/529204 (2008).

32 Yeaman, M. et al. in 52nd Interscience Conference on Antimicrobial Agents and Chemotherapy.

33 Yeaman, M. R. et al. in Gordon Research Conference on Staphylococcal Diseases.

34 Paczosa, M. K. & Mecsas, J. Klebsiella pneumoniae: Going on the Offense with a Strong Defense. Microbiol Mol Biol Rev 80, 629–661, DOI:10.1128/MMBR.00078-15 (2016).

35 Curtin, F. et al. GNbAC1, a humanized monoclonal antibody against the envelope protein of multiple sclerosis-associated endogenous retrovirus: a first-in-humans randomized clinical study. Clin Ther 34, 2268–2278, DOI:S0149-2918(12)00649-2 [pii] 10.1016/j.clinthera.2012.11.006.

36 Xu, L. et al. Pharmacokinetics of ranibizumab in patients with neovascular age-related macular degeneration: a population approach. Invest Ophthalmol Vis Sci 54, 1616–1624, DOI:iovs.12-10260 [pii] 10.1167/iovs.12-10260.

37 Gebremariam, T. et al. CotH3 mediates fungal invasion of host cells during mucormycosis. J Clin Invest 124, 237–250, DOI:10.1172/JCI71349 (2014).

